# Using balances to engineer features for the classification of health biomarkers: a new approach to balance selection

**DOI:** 10.1101/600122

**Authors:** Thomas P. Quinn, Ionas Erb

## Abstract

Since the turn of the century, technological advances have made it possible to obtain a molecular profile of any tissue in a cost-effective manner. Among these advances include sophisticated high-throughput assays that measure the relative abundance of microorganisms, RNA molecules, and metabolites. While these data are most often collected to gain new insights into biological systems, they can also be used as biomarkers to create clinically useful diagnostic classifiers. How best to classify high-dimensional “-omics” data remains an area of active research. However, few explicitly model the relative nature of these data, and instead rely on cumbersome normalizations which often invoke untestable assumptions. This report (a) emphasizes the relative nature of health biomarkers, (b) discusses the literature surrounding the classification of relative data, and (c) benchmarks how different transformations perform across multiple biomarker types. In doing so, this report explores how one could use balances to engineer features prior to classification, and proposes a simple procedure, called discriminative balance analysis, to select discriminative 2- and 3-part balances.

## 1 Introduction

Next-generation sequencing (NGS) technology is routinely used to quantify the presence of bacterial or gene species from environmental and biological samples. Hyphenated chromatographic assays like liquid chromatography-mass spectrometry (LC-MS) are used to quantify the presence of proteins, lipids, or metabolites. NGS and LC-MS both generate high-dimensional data that are used as health biomarkers to predict and surveil disease [28]. However, because NGS and LC-MS measure abundance by sampling from the total population, the total number of molecules recorded for a sample is arbitrary, thereby making these data compositional [26, 14, 30, 21, 22, 20, 44]. Others have already demonstrated that compositionality confounds the routine application of univariate [6], correlation [40], and distance [2] measures. Since machine learning pipelines often rely on these measures, compositionality may impact the accuracy of classifiers trained on these data [14, 24].

Compositional data analyses tend to have one of three flavours. First, the “simple” log-ratio approach uses a single reference to re-cast the data. Most commonly, the reference is the per-sample geometric mean (centered log-ratio [clr] transformation) or a single component (additive log-ratio [alr] transformation), but the geometric mean of inter-quartile range components [55] and of nonzero components [31] have also been proposed. After transformation, the analysis then proceeds as if the data were absolute, but with a caveat: the interpretation of the results depends on the reference used. Second, the “pragmatic approach” analyzes pairwise log-ratios directly, which has been used to score important genes [52] and gene pairs [23, 13], and to reduce the dimensionality of the data [23]. This approach makes sense when the ratios themselves have some importance to the analyst. However, it presents a clear problem for the classification of high-dimensional data: ratios “explode” feature space from *p* = *D* features to *p* = *D*(*D* − 1)/2 (pairs of) features, compounding the *p* >> *N* problem. Third, the “coordinate approach” uses an orthonormal basis to transform *D* components into *D* – 1 values via an isometric log-ratio (ilr) transformation [12]. When a serial binary partition (SBP) matrix is used to obtain the orthonormal basis, this approach yields a set of “balances”, each of which describe a log-contrast between two sets of components [11, 49, 38]. Since the basis vectors used to make balances are associated with successive bipartitions of the original feature set, balances can be more interpretable than general log-contrasts, while still having the formal appeal of the ilr transformation (i.e., orthogonality of the basis vectors and a full-rank covariance matrix) [11, 7]. However, the utility of balances depends on having a desirable SBP matrix, which must be manually curated or procedurally generated. One popular SBP, easily approximated via a simple heuristic, decomposes the variance such that the first balance explains the most variance, the second balance the second most, and so on [39, 32]. In microbiome research, authors have proposed using mean pH [35] and phylogenetics [48, 53] to construct an SBP.

Many publications have applied *unsupervised* statistical learning to compositional data. This includes the application of principal components analysis (PCA) [1], PCA biplots [4], and cluster analysis [33]. For PCA analysis, a clr-transformation is often used, although clr data do not have a full rank [15]. Instead, a robust PCA could be applied to ilr-transformed data, with the loadings back-transformed for interpretation [15].For cluster analysis, the Aitchison distance is preferred over the Euclidean distance, because only the former is sub-compositionally dominant [3, 33, 2]. Recently, Martin-Fernandez et al. have applied self-organized maps, a non-linear dimension reduction method, to ilr-transformed data [34], while Martino et al. have extended compositional PCA to zero-laden microbiome data [31]. Several studies have also applied *supervised* statistical learning to compositional data. Aitchison trained linear discriminant analysis (LDA) models on alr-transformed data [1], as have others [9] (though LDA is now usually applied to ilr-transformed data [9, 8]). Generalized linear models, including logistic regression (LR), have also been used to classify composition data [50, 8]. However, both LDA and LR require at least as many samples as features, making them inappropriate for high-dimensional health biomarker data (though this limitation is mitigated by regularization, used by [51] and [29] to classify compositions). Partial least squares (PLS), also suitable for high-dimensional data, has been applied to clr-transformed data to predict continuous outcomes [25], while PLS discriminant analysis (PLS-DA) has been used to classify clr-transformed [18] and ilr-transformed [27] data. In microbiome research, a step-wise algorithm, implemented as selbal, was proposed to identify a single balance that performs well in classification and regression tasks [46]. This latter work highlights an advantage to balances: although alr-, clr-, and ilr-transformations can facilitate statistical learning, balances can engineer the feature space into a single interpretable biomarker via *balance selection*.

How best to classify high-dimensional compositional data remains an open question. We are not aware of any work that benchmarks compositional data transformations as they pertain to the classification of high-dimensional compositional data. In this study, we employ a statistically robust cross-validation scheme to evaluate how well regularized LR classifies health-related binary outcomes on 13 compositional data sets. Specifically, we benchmark performance using features obtained from raw proportions, clr-transformed data, balances, and selected balances. Our results show that the centered log-ratio transformation, and all four balance procedures, outperform raw proportions for the classification of health biomarker data. We also propose a new balance selection procedure, called discriminatory balance analysis, that offers a computationally efficient way to select important 2- and 3-part balances. These discriminant balances reduce the feature space and improve the interpretability, without sacrificing classifier performance. In doing so, they also outperform a recently published balance selection method, selbal, in terms of run-time and classification accuracy.

## 2 Methods

### 2.1 Data acquisition

We acquired data from 4 principal sources. Two gut microbiome data sets (originally published in [19] and [37]) were acquired already quantified from the selbal package [46]. Two additional gut microbiome data sets (originally published in [47] and [5]) were acquired already quantified from the supplement of Duvallet et al. [10]. A fifth gut microbiome data set was acquired already quantified from the supplement of Franzosa et al. [16].

The Schubert et al. data contained 3-classes comparing hospital-acquired diarrhea (HAD) with community-acquired diarrhea (CAD) and healthy controls. This data set was used in two tests: HAD vs. CAD and HAD vs. HC. The Buxter et al. data contained 3-classes comparing colorectal cancer (CRC) with adenoma (AC) and healthy controls (HC). This data set was also used in two tests: CRC vs. AC and CRC vs. HC. The Franzosa et al. data contained 3-classes comparing Crohn’s disease (CD) and ulcerative colitis (UC) with healthy controls (HC). This data set was also used in two tests: (CD & UC) vs. HC and CD vs. UC. Franzosa et al. also published gut metabolomic data for the same samples. These data were used for an additional two tests that parallel the gut microbiome tests.

A sixth data set was acquired from The Cancer Genome Atlas [54], and contains microRNA expression for primary breast cancer (BRCA) samples and healthy controls (HC). We further labelled the BRCA samples using PAM50 sub-types retrieved from the supplement of [36]. PAM50 uses a gene expression signature to assign an intrinsic sub-type to the primary breast cancer sample: sub-types include luminal A, luminal B, HER2-enriched, Basal, and Normal-like. These data were used in three tests: any BRCA vs. HC, HER2+ vs. all other BRCA, and LumA-BRCA vs. LumB-BRCA.

### 2.2 Feature extraction and zero handling

Before training any models, features with too few counts were removed from the data. For the metabolomic and microRNA data sets, only features within the top decile of total abundance were included (this was done to reduce the feature space so that selbal became computationally tractable). For all data sets, features that contained zeros in more than 90% of samples were excluded (this was done to remove biomarkers that are not reliably present in the data). Finally, zeros were replaced using a simple multiplicative replacement strategy via the zCompositions package (this was done because the Bayesian replacement strategy fails for heavily zero-laden data). Table 1 summarizes the tests used in this study.

**Table 1:**
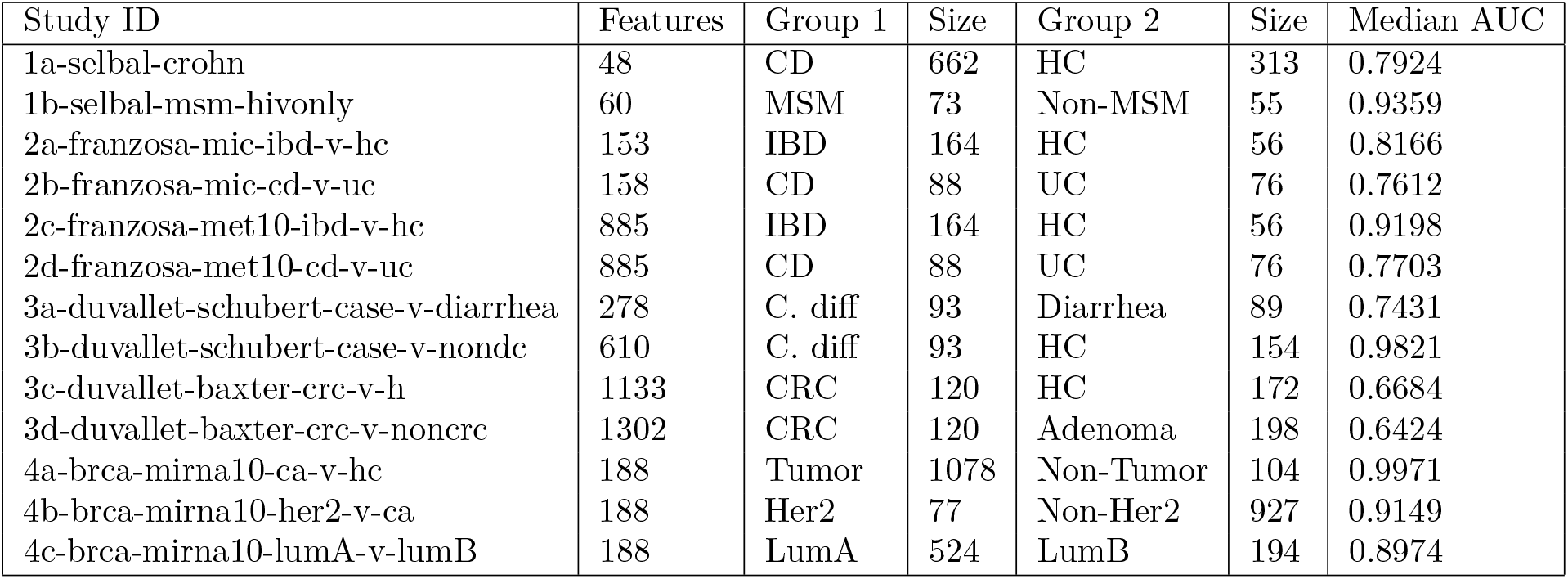
This table describes the data used to benchmark data transformation and balance selection methods. Acronyms: CD Crohn’s disease; HC healthy control; MSM men who have sex with men; UC ulcerative colitis; IBD inflammatory bowel disease; CRC colorectal cancer.

### 2.3 Data transformation

Let us consider the data matrix *x_ij_* which describes the relative abundance of *j* ∈ {1,…, *D*} components (as features) across *i* ∈ {1,…,*N*} compositions (as samples). Since the data studied are compositional, they can be expressed as a sub-composition of parts of the whole. The closure operation expresses the data so that the measurements for each sample sum to 1 (i.e., as proportions). The closed data are benchmarked in this study as the point of reference:

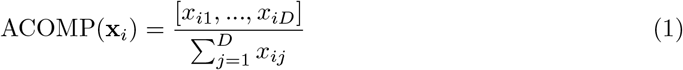

We also benchmark the popular centered log-ratio transformation (**clr**):

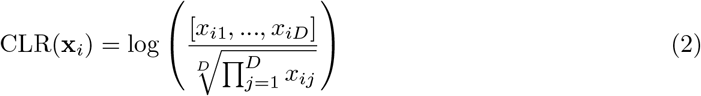

We also use the isometric log-ratio transformation (ilr) to construct balances from a serial binary partition (**SBP**) matrix describing *z* ∈ {1,…, *D* − 1} log-contrasts between *j* ∈ {1,…, *D*} parts. A log-contrast of a *D*-part composition is a log-linear combination *a*_1_ log *x*_*i*1_ + ⋯ + *a_D_* log *x_iD_* with the constraint that 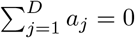. This constraint ensures scale invariance of the combination, i.e., normalization factors of **x**_*i*_ cancel.

Balances are constructed from special log-contrasts with 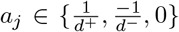 (where *d*^+^ and *d*^−^ refer to the number of positive and negative entries in a column of the SBP matrix). Such log-contrasts have the form 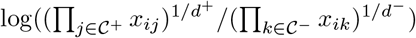 where 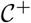 and 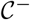 are the sets of indices *j* for which 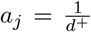 and 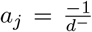, respectively. The special form of these log-contrasts allows for a representation in form of a bi-partition. It is helpful to think of an SBP as a dendrogram tree, from which the terms *a_j_* can be derived (see Figure 1 for an example SBP). A balance value is now computed for each sample *i* and each log-contrast *z*:

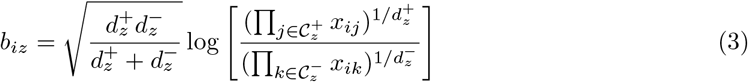

**Figure 1:**
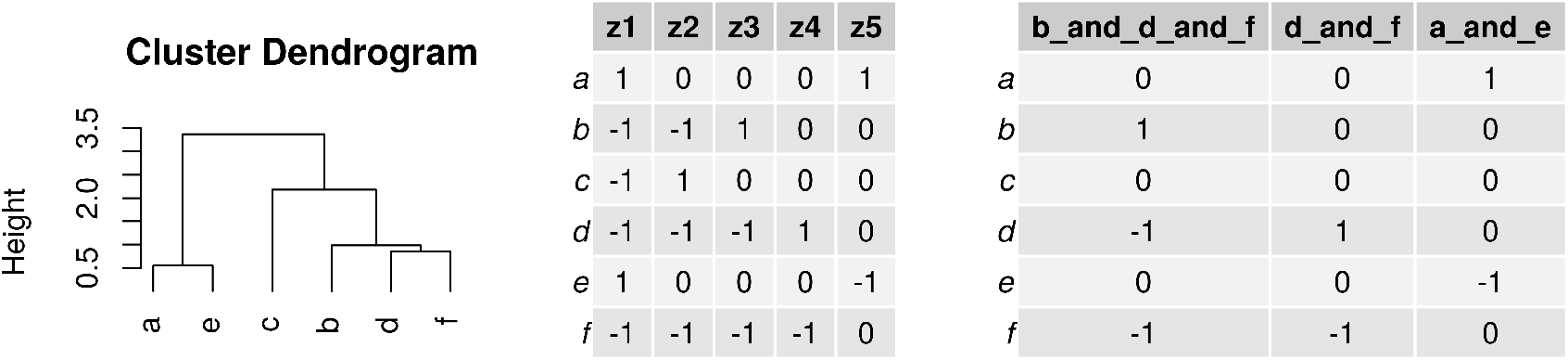
This figure shows how a balance dendrogram relates to a serial binary partition (SBP) matrix. The left panel shows a dendrogram clustering the similarity between 6 components, where the first branch in the dendrogram refers to the first balance (i.e., [a & e] vs. [c & b & d & f]). The middle panel shows the corresponding SBP with 5 balances (columns) and the components involved in each log-contrast (rows). The right panel shows the “distal” 2- and 3-part balances.

The prefactor standardizes the log-contrasts such that 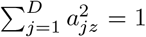 and makes them orthogonal in the sense that 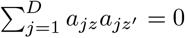 for all *z* ≠ *z*′.

### 2.4 The serial binary partition matrix

We benchmark four procedures for generating an SBP. In **PBA**, we approximate a set of principal balances by hierarchically clustering the log-ratio variance matrix, **T**, describing the relationship between any two variables *j* and *j*\* (see [39]):

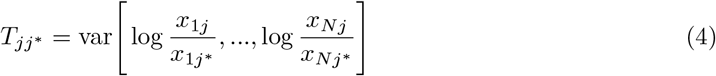

Principal balances are analogous to principal components in that the first balance explains the most variance, the second balance the second most variance, and so on. Note that **PBA** only approximates principal balances.

In **ABA**, we hierarchically cluster a new dissimilarity measure defined as the difference of the log-ratio variance matrix from the maximum log-ratio variance score: max(**T**) − *T*_*jj*\*_. In **RBA**, we generate random SBPs using a custom algorithm. In DBA, we generate an SBP that maximizes the discriminative potential of the distal branches. This is done by hierarchically clustering the differential proportionality matrix, Θ, describing the relative contribution of the within-group logratio variances (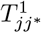 and 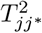) to the total log-ratio variance (see [13, 43]):

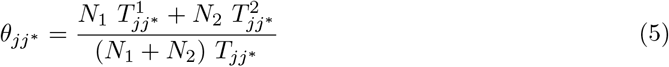

for groups sized *N*_1_ and *N*_2_. This matrix ranges from [0, 1] where 0 indicates that the two features have a maximally large difference in log-ratio means between the two group. Unlike the other SBP methods, the **DBA** method is supervised.

Note that the SBP is always constructed using the *training set only*. The balance “rule” is then applied to the validation set prior to model deployment. All SBP procedures are implemented in the `balance` package with the functions `sbp.fromPBA`, `sbp.fromABA`, `sbp.fromRandom`, and `sbp.fromPropd`, respectively [42]. Differential proportionality analysis is implemented in the `propr` package [45] with the function `propd`.

### 2.5 Classification pipeline

In order to get a robust measure of performance, we repeat model training on 50 training sets randomly sampled from the data (with 33% set aside as a validation set). For each training set, we (1) transform features as described above, (2) train a model on the transformed features, (3) deploy the model on the withheld validation set, and (4) calculate the area under the receiver operating curve (AUC). AUC is used because it is more informative than accuracy when classes are imbalanced. Model splitting, transformation, training, and prediction are all handled by the high-throughput classification software `exprso` [41]. By repeating this procedure 50 times, we can calculate the median performance and its range.

When using `selbal`, a generalized linear model is trained on a single balance (as done in [46]). For all other transformations, a least absolute shrinkage and selection operator (LASSO) model is used to select features and fit the data simultaneously (via the `glmnet` package [17]). When using LASSO, λ is chosen procedurally by measuring 5-fold training set cross-validation accuracy over the series `exp(seq(log(0.001), log(5)`, `length.out=100))`, with the best λ selected automatically by `cv.glmnet`.

## 3 Results and Discussion

### 3.1 Choice in log-ratio transformation does not impact performance

Figure 2 shows the validation set AUCs for binary classifiers trained on 13 data sets. In general, we see that the centered log-ratio transformation (CLR) and balance procedures (PBA, ABA, RBA, DBA) perform comparably. Although they all tend to outperform proportions (ACOMP), the proportions were more discriminative than the CLR for a few tests. This might occur when the closure bias itself confounds the predicted outcome.

**Figure 2:**
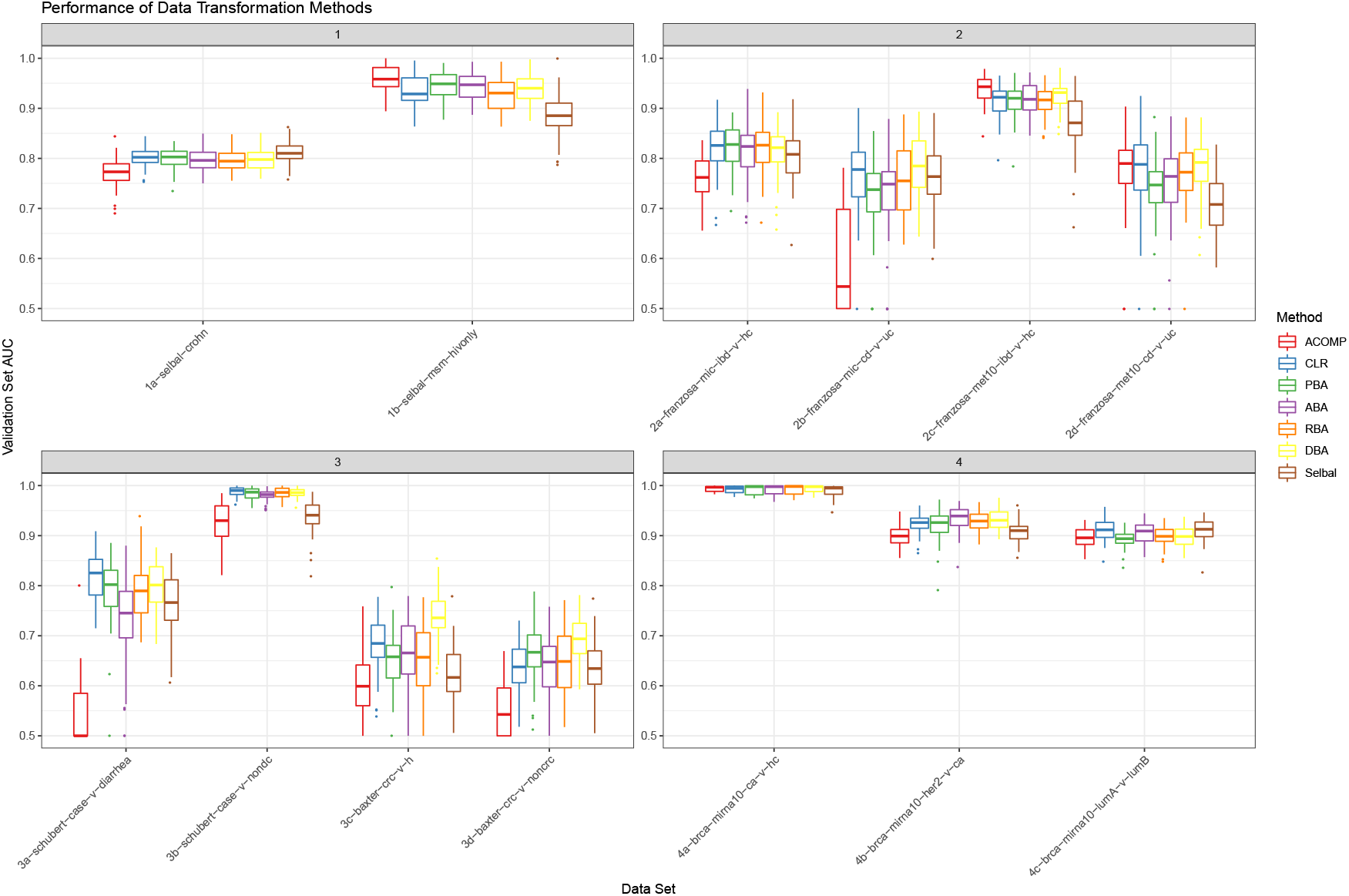
This figure shows the distribution of validation set AUCs (y-axis) for classifiers trained on closed or transformed data (x-axis). Each validation set AUC describes a unique random training and validation set split. All classifiers are regularized logistic regression models, with λ tuned by training set cross-validation. Acronyms: ACOMP closed proportions; CLR centered log-ratio transformed data; PBA principal balances; ABA anti-principal balances; RBA random balances; DBA discriminative balances.

Table 2 shows the median of the difference between data transformations (as computed with pairwise Wilcoxon Rank Sum tests across all 13 tests). Here, we see that every transformation performs better than proportions. We also note that all balance procedures tend to perform equally well, though DBA balances perform marginally better. Although `selbal` posts an impressive accuracy for only using a single balance, it is less accurate than using a set of all balances.

**Table 2:** This table shows the median of the difference in performance between data transformation methods. Confidence intervals computed using pairwise Wilcoxon Rank Sum tests applied to 50 re-samplings of 13 data sets. Acronyms: ACOMP closed proportions; CLR centered log-ratio transformed data; PBA principal balances; ABA anti-principal balances; RBA random balances; DBA discriminative balances.

|  | Selbal vs. | PBA vs. | ABA vs. | RBA vs. | DBA vs. | ACOMP vs. | CLR vs. |
| --- | --- | --- | --- | --- | --- | --- | --- |
| Selbal | — | 0.0054 to 0.0324 | 0.0047 to 0.0318 | 0.0074 to 0.0339 | 0.019 to 0.045 | -0.0290 to 0.0013 | 0.014 to 0.040 |
| PBA | -0.0324 to -0.0054 | — | -0.013 to 0.011 | -0.0097 to 0.0138 | 0.0013 to 0.0248 | -0.048 to -0.016 | -0.003 to 0.020 |
| ABA | -0.0318 to -0.0047 | -0.011 to 0.013 | — | -0.0092 to 0.0148 | 0.0018 to 0.0253 | -0.045 to -0.015 | -0.0026 to 0.0204 |
| RBA | -0.0339 to -0.0074 | -0.0138 to 0.0097 | -0.0148 to 0.0092 | — | -0.00061 to 0.02223 | -0.048 to -0.017 | -0.0048 to 0.0177 |
| DBA | -0.045 to -0.019 | -0.0248 to -0.0013 | -0.0253 to -0.0018 | -0.02223 to 0.00061 | — | -0.060 to -0.029 | -0.0144 to 0.0065 |
| ACOMP | -0.0013 to 0.0290 | 0.016 to 0.048 | 0.015 to 0.045 | 0.017 to 0.048 | 0.029 to 0.060 | — | 0.024 to 0.054 |
| CLR | -0.040 to -0.014 | -0.020 to 0.003 | -0.0204 to 0.0026 | -0.0177 to 0.0048 | -0.0065 to 0.0144 | -0.054 to -0.024 | — |

### 3.2 DBA method selects predictive balances

An advantage of using regularized logistic regression is that the model weights can be interpreted as a measure of feature importance. Even though the CLR and balances perform equally well, they imply different interpretations. While the CLR data do have a feature representing each component, the regularized weights do not describe the importance of *that component*. Rather, the CLR weights describe the importance of *that variable relative to the sample mean*. On the other hand, balances measure the log-contrast between sets of components. Thus, the balance weights describe the importance of those components directly.

For high-dimensional data, it can be challenging to interpret large balances. For example, the base of an SBP always contains one balance that is comprised of all variables. It may not be helpful in understanding the outcome to know that a log-contrast involving all components is discriminative. On the other hand, smaller balances (i.e., those involving fewer components) might have a clearer meaning to the analyst. Here, we propose a new procedure, called *discriminative balance analysis* (DBA), to generate an SBP that makes the smallest balances most discriminative. This procedure can be used to engineer and select important balances prior to model building. Since the selected balances contain few parts, they are more easily interpreted.

Conceptualizing the SBP as a tree, the largest balances are the “trunk” and the smallest balances are the “leaves” (see Figure 1). Since the basis is orthonormal, we can consider segments of the tree in the isolation of others. This also makes sure that the selected balances do not contain redundant information. Figure 3 shows classification AUC when only using the “distal leaf” balances (i.e., those with 2- or 3- parts). In principal balances analysis (PBA), the “trunk” contains the most variance, and the leaves the least. As expected, the distal PBA balances perform poorly. In anti-principal balance analysis (ABA), the “trunk” contains the least variance, and the leaves the most. As expected, the distal ABA balances outperform the distal PBA balances. In discriminative balance analysis (DBA), the “trunk” is least discriminative, and the leaves the most. As expected, the distal DBA balances outperform both the PBA and ABA balances. Indeed, since DBA places the most discriminative balances distally, the distal DBA balances perform as well as all DBA balances (see Table 3 for 95% confidence interval). In random balance analysis (RBA), balances are random, so the leaves might be discriminative by chance. As expected, the distal RBA balances have an average performance.

**Figure 3:**
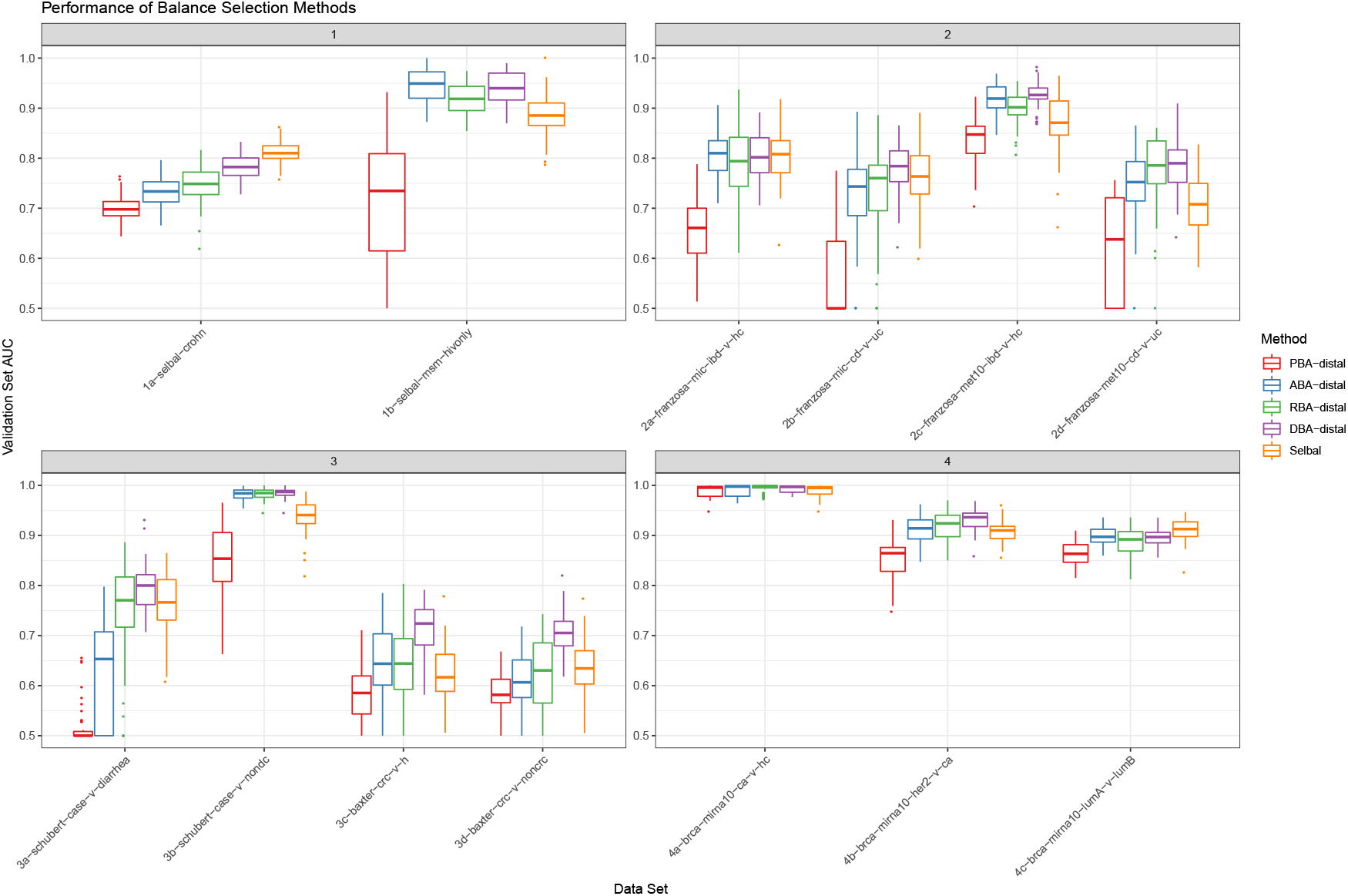
This figure shows the distribution of validation set AUCs (y-axis) for classifiers trained on selected balances (x-axis). Each validation set AUC describes a unique random training and validation set split. All classifiers are regularized logistic regression models, with λ tuned by training set cross-validation. Acronyms: PBA principal balances; ABA anti-principal balances; RBA random balances; DBA discriminative balances. The appendix “-distal” indicates that only the 2-part and 3-part balances were used as features.

**Table 3:** This table shows the median of the difference in performance between balance selection methods. Confidence intervals computed using pairwise Wilcoxon Rank Sum tests applied to 50 re-samplings of 13 data sets. Acronyms: PBA principal balances; ABA anti-principal balances; RBA random balances; DBA discriminative balances. The appendix “-distal” indicates that only the 2-part and 3-part balances were used as features.

|  | Selbal vs. | PBA-distal vs. | ABA-distal vs. | RBA-distal vs. | DBA-distal vs. | DBA vs. |
| --- | --- | --- | --- | --- | --- | --- |
| Selbal | — | -0.125 to -0.091 | -0.012 to 0.016 | -0.0066 to 0.0203 | 0.016 to 0.042 | 0.019 to 0.045 |
| PBA-distal | 0.091 to 0.125 | — | 0.087 to 0.122 | 0.095 to 0.131 | 0.12 to 0.16 | 0.13 to 0.16 |
| ABA-distal | -0.016 to 0.012 | -0.122 to -0.087 | — | -0.0082 to 0.0182 | 0.014 to 0.040 | 0.017 to 0.044 |
| RBA-distal | -0.0203 to 0.0066 | -0.131 to -0.095 | -0.0182 to 0.0082 | — | 0.0082 to 0.0345 | 0.012 to 0.038 |
| DBA-distal | -0.042 to -0.016 | -0.16 to -0.12 | -0.040 to -0.014 | -0.0345 to -0.0082 | — | -0.006 to 0.014 |
| DBA | -0.045 to -0.019 | -0.16 to -0.13 | -0.044 to -0.017 | -0.038 to -0.012 | -0.014 to 0.006 | — |

The weights of DBA balances can be interpreted (and visualized) in an intuitive way. The 2-part balances can be visualized as a log-ratio, while the 3-part balances can be visualized with a ternary diagram or as a log-contrast. In Figure 4, we compare the most important distal DBA balances (left panel) with the single discriminative balance found by selbal (right panel). We see that many of the same variables are represented in both sets. However, DBA expresses the important variables via 2- and 3-part subsets that are, by definition of the SBP, grouped to be maximally discriminative. In the left panel, we see that balances with large regularized weights (top-left) have log-contrast scores that differentiate the groups (bottom-left). Though selbal performs remarkably well in its ability to select a single discriminative balance, our results suggest that the distal DBA method outperforms selbal by ~1-4% AUC (see Table 3). Moreover, the distal DBA method is an order of magnitude faster than selbal, the latter of which must try multiple component combinations before finding the best log-contrast (25 minutes vs. 15 seconds for 1000 features).

**Figure 4:**
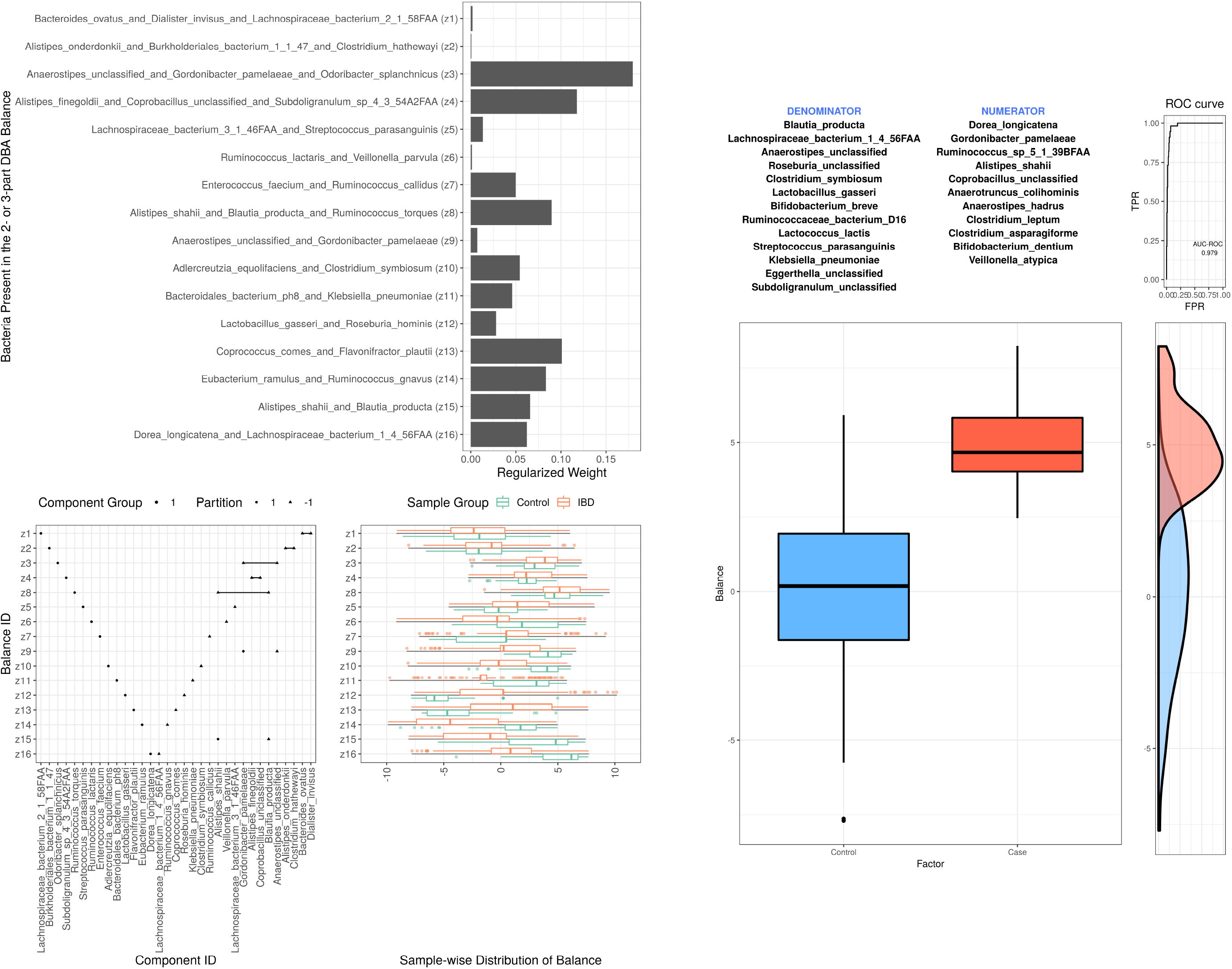
This figure shows the most important distal DBA balances (top-left panel), their composition and log-contrast scores (bottom-left panel), and the single discriminative balance found by `selbal` (right panel). All panels generated using the `2a-franzosa-mic-ibd-v-hc` data set.

### 3.3 DBA as a discriminant ordination

By using an orthonormal basis, balances decompose the variance into orthogonal parts. For a binary outcome, this allows us to not only calculate the amount of variance contained in each discriminative balance, but also to break down the contained variance into its *between-group* and *within-group* fractions (as done by an ANOVA). The left panel of Figure 5 shows that a large fraction of the (log-ratio) variance contained in the distal DBA balances is *between-group variance*. This is because the clustering of components using *θ*_*jj*\*_ as a distance will group together those components whose pairwise log-ratios describe only a small fraction of the within-group variance (i.e., a large fraction of between-group variance). Although we have already shown that the distal DBA balances are discriminative, the orthonormality of their underlying basis also allows us to use DBA to project a discriminant ordination of the data. In other words, we can visualize the data along mutually exclusive discriminant axes (similar to the axes in a discriminant analysis decomposing the variance between group means; however, for two groups, this would only give a single axis). The right panel of Figure 5 shows good class separation using only 3 balances (each of which is actually a simple log-ratio). From the left panel, we know that these three axes contain 4.3% of the total variance, and could likewise calculate that they contain 13.8% of the total between-group variance. Meanwhile, the DBA-selected balances together account for 90.4% of the total between-group variance. Yet, by restricting the ordination to the distal DBA balances, we ensure that each one of these discriminant axes is fully interpretable, having no more than 3 parts. On the other hand, if the analyst cared less about interpretation and more about maximizing contained between-group variance, they could do a clustering using as distance 1 − *θ*_*jj*\*_ and instead project the largest balances thus obtained (in direct analogy to the principal balances heuristic described above).

**Figure 5:**
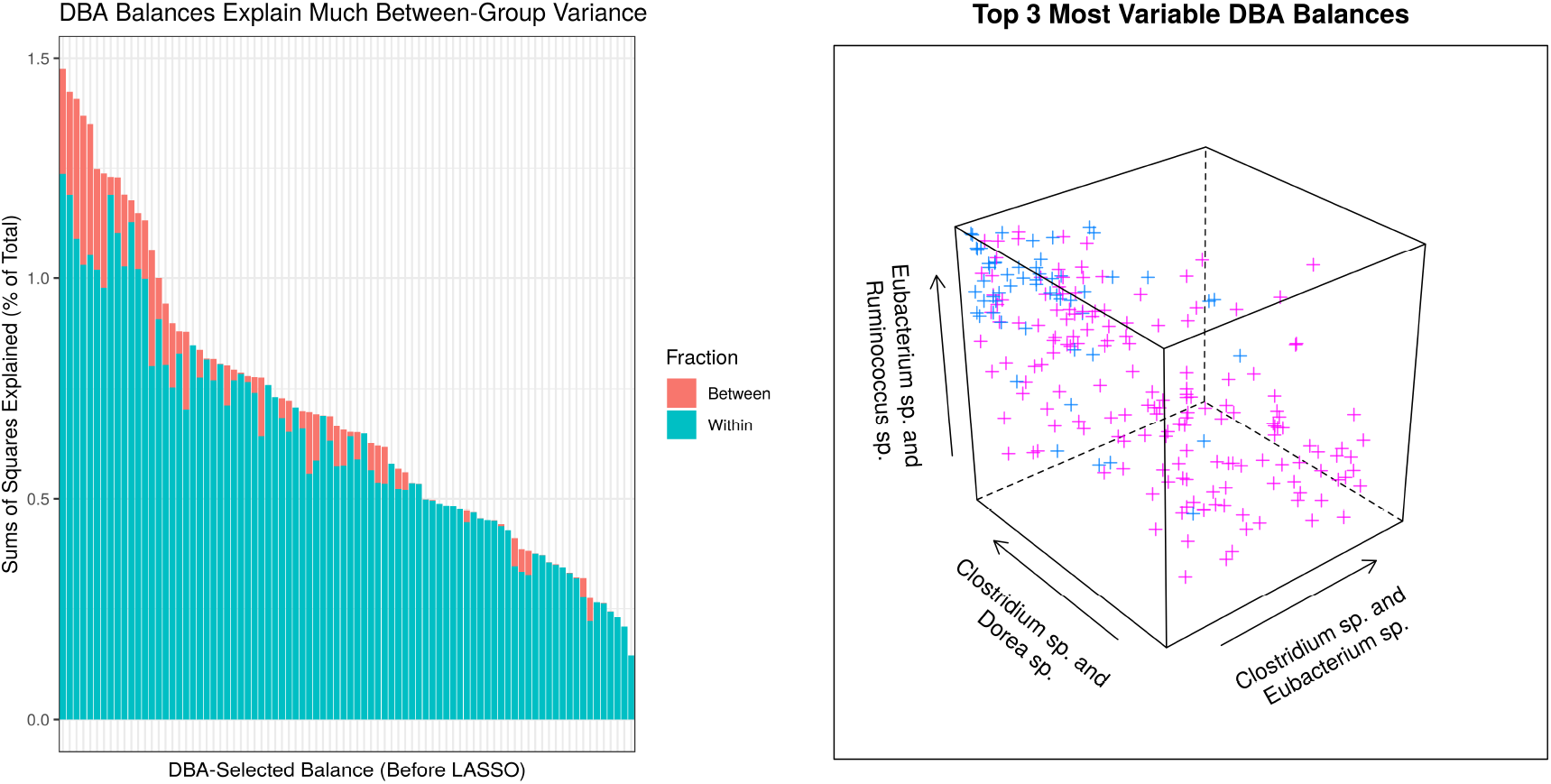
This figure shows the amount of variance (as a percent of the total) contained in each distal DBA balance (left panel), placed alongside a projection of the data across the top 3 most variable distal DBA balances (right panel). Good class separation is achieved using only 3 balances (each of which is proportional to a simple log-ratio). All panels generated using the `2a-franzosa-mic-ibd-v-hc data set`.

## 4 Summary

This work benchmarks the performance of regularized logistic regression classifiers across 13 high-dimensional health biomarker data sets. Our results show that, on average, the centered log-ratio and balances both outperform raw proportions in classification tasks. We find that the serial binary partition (SBP) matrix used to generate the balances does not impact performance. However, the choice in SBP changes *which* balances are important for classification. In this report, we introduce a new SBP procedure that makes the most discriminative balances the smallest. This procedure, called *discriminative balance analysis,* offers a computationally efficient way to select important 2- and 3-part balances. These discriminant balances reduce the feature space and improve the interpretability, without sacrificing classifier performance. In doing so, they also outperform a recently published balance selection method, selbal, in terms of run-time and classification accuracy.

## 5 Declarations

### 5.1 Ethics approval and consent to participate

Not applicable.

### 5.2 Consent for publication

Not applicable.

### 5.3 Availability of data and material

All methods are available through open source software maintained by the authors.

### 5.4 Competing interests

No authors have competing interests.

### 5.5 Funding

Not applicable.

### 5.6 Authors’ contributions

TPQ implemented the procedures, performed the analyses, and drafted the manuscript. IE derived the differential proportionality metric, contributed code, and expanded the manuscript. Both authors conceptualized the thesis and approved the final manuscript.

## 6 Acknowledgements

TPQ thanks the authors of selbal for inspiring this work. TPQ thanks Samuel C. Lee for his help with retrieving the TCGA data and the PAM50 labels. IE thanks Cedric Notredame for support.

